# Conserved intramolecular networks in GDAP1 are closely connected to CMT-linked mutations and protein stability

**DOI:** 10.1101/2022.09.09.507262

**Authors:** Aleksi Sutinen, Dirk Paffenholz, Giang Thi Tuyet Nguyen, Salla Ruskamo, Andrew E. Torda, Petri Kursula

## Abstract

Charcot-Marie-Tooth disease (CMT) is the most common inherited peripheral polyneuropathy in humans, and its subtypes are linked to mutations in dozens of different genes, including the gene coding for ganglioside-induced differentiation-associated protein 1 (GDAP1). The main GDAP1-linked CMT subtypes are the demyelinating CMT4A and the axonal CMT2K. Over a hundred different missense CMT mutations in the *GDAP1* gene have been reported. However, despite implications for mitochondrial fission and fusion, cytoskeletal interactions, and response to reactive oxygen species, the etiology of GDAP1-linked CMT is poorly understood at the protein level. Based on earlier structural data, CMT-linked mutations could affect intramolecular interaction networks within the GDAP1 protein. We carried out structural and biophysical analyses on several CMT-linked GDAP1 protein variants and describe new crystal structures of the autosomal recessive R120Q and the autosomal dominant A247V and R282H GDAP1 variants. These mutations reside in the structurally central helices α3, α7, and α8. In addition, solution properties of the CMT mutants R161H, H256R, R310Q, and R310W were analysed. All disease variant proteins retain close to normal structure and solution behaviour. All mutations, apart from those affecting Arg310 outside the folded GDAP1 core domain, decreased thermal stability. In addition, a bioinformatics analysis was carried out to shed light on the conservation and evolution of GDAP1, which is an outlier member of the GST superfamily. Many CMT mutation sites are highlighted in the bioinformatics analyses focusing on sequence conservation and entropy, and the analyses support the outlier nature of GDAP1 in the GST superfamily. A central role for the α6-α7 loop, within a conserved interaction network, is identified for GDAP1 protein stability. To conclude, we have expanded the structural analysis on GDAP1, strengthening the hypothesis that alterations in conserved intramolecular interactions may alter GDAP1 stability and function, eventually leading to mitochondrial dysfunction, impaired protein-protein interactions, and neuronal degeneration.

## INTRODUCTION

The demand for sufficient energy supply *via* the aerobic process is elevated in neurons compared to other organelles and tissues, such as muscles [1]. Mitochondria are responsible for cellular respiration and linked to Ca^2+^ signalling and reactive oxygen species metabolism [2–5]. Since neurons depend on aerobic energy, their demand for oxidative phosphorylation is high, and 20% of the net oxygen consumed by the body is used for oxidative phosphorylation in neurons. Therefore, neurons are sensitive to alterations in mitochondrial function, and disruptions in mitochondrial dynamics can have severe consequences on neuronal functions.

Mitochondria are not isolated organelles, but interact with other cellular compartments, such as the endoplasmic reticulum, lysosomes, and peroxisomes, exchanging metabolites [2, 6, 7]. Mitochondria are renewed *via* fission and fusion, which are driven by proteins on the mitochondrial outer membrane (MOM), such as mitofusin 1 and 2 (MFN1/2), and the mitochondrial inner membrane (MIM), such as OPA1 and FIS1. Auxiliary proteins may bind to either MOM or MIM to enhance the process. The ganglioside-induced differentiation-associated protein 1 (GDAP1) is an integral MOM protein, proposed to have an auxiliary role in mitochondrial fission and fusion [8], possibly *via* redox-dependent interactions with cytoskeletal components [9]. However, the molecular basis of GDAP1 function and its exact relation to disease are currently not known.

Structurally, GDAP1 resembles glutathione *S*-transferases (GST), and it contains unique flexible loops [10, 11]. GDAP1 is constructed of two GST-like domains in the N and C terminus, followed by a transmembrane helix, which anchors the protein into the MOM. Structural data have shown a covalently bound dimer interface in GDAP1 [11, 12], and while dimerization is a common feature in catalytic GSTs [13], the GDAP1 dimer is formed differently [11]. Enzymatic activity of GDAP1 has not been convincingly demonstrated, nor has any substrate been identified *in vivo.* The members of the GST superfamily have similar fold properties, whereas distinct differences and low sequence conservation result in a diverse group of substrates; hence, it is possible that GDAP1 is an enzyme, but the substrate and reaction mechanism remain unidentified.

Increasing numbers of genes related to mitochondrial functions have been linked to neuropathophysiological conditions. Inherited polyneuropathies are a genetically and clinically diverse group of neurodegenerative diseases, which affect the outer motor and sensory neurons in the peripheral nervous system (PNS) [14, 15], the most common being Charcot-Marie-Tooth disease (CMT). Clinical profiling divides CMT into three classes: demyelinating, axonal, and intermediate [16, 17]. The phenotype often implies insufficient mitochondrial fission and fusion, and mitochondria appear fragmented and elongated [18]. The etiology of CMT is linked to the hereditary pattern, whereby the autosomal recessive form has an earlier onset and more severe symptoms than the autosomal dominant form [19–21]. In the case of GDAP1, both autosomal dominant (axonal type CMT2) and recessive (demyelinating type CMT4) modes of inheritance are found. The severity is often correlated with the location of the causative mutation in the protein. The *GDAP1* gene is one of the most common missense mutation targets linked to CMT [8, 22, 23]. GDAP1 is ubiquitously expressed in tissues, but most of the expression is confined to neuronal tissues [8, 24]. The most accurate structural data thus far cover the dimeric core GST-like domain of human GDAP1, including the GDAP1-specific insertion [11]. In addition, a structure of a construct missing the large GDAP1-specific insertion is available in monomeric form [10]. In full-length GDAP1, an amphipathic extension – originally termed the hydrophobic domain – links the transmembrane helix to the GST-like domain.

GSTs often contribute to mechanisms of neurodegenerative disease [25, 26]. GST superfamily members function in prokaryotic and eukaryotic metabolism by utilizing reduced glutathione to catalyse a range of chemically diverse reactions. In comparison to other enzyme superfamilies, GSTs are unique in that sequence conservation appears to be driven by fold stability instead of catalytic features, as reflected in the broad spectrum of GST substrates [27, 28]. Using X-ray crystallography and complementary biophysical and computational techniques, we carried out structural analysis on selected GDAP1 mutants linked to CMT. We also analysed GDAP1 sequence conservation to investigate its GST-linked ancestry and to get clues into its molecular function and the relationship between conserved residue interaction networks and disease mutations.

## MATERIALS AND METHODS

### Recombinant protein production and purification

The GDAP1Δ303-358 and GDAP1Δ319-358 constructs, with an N-terminal His_6_ tag and a Tobacco Etch Virus (TEV) protease digestion site, for producing soluble recombinant human GDAP1 in *E. coli*, have been described [11]. The point mutations R120Q, R161H, A247V, H256R, and R282H were generated in GDAP1Δ303-358, and the mutations R310Q and R310W in GDAP1Δ319-358, by a site-directed mutagenesis protocol with Pfu polymerase. All constructs were verified by DNA sequencing.

Recombinant GDAP1 variants were expressed in *E. coli* BL21(DE3) using autoinduction [29], and purified as described [11]. Briefly, GDAP1 was separated from the lysate by Ni^2+^-NTA chromatography, and the affinity tag was cleaved using TEV protease. Another Ni^2+^-NTA affinity step removed the tag and TEV protease. Size exclusion chromatography (SEC) was performed on a Superdex 75 10/300 GL increase column (Cytiva) using 25 mM HEPES (pH 7.5) and 300 mM NaCl (SEC buffer) as mobile phase. SEC peak fractions were analysed with SDS-PAGE and concentrated by centrifugal ultrafiltration.

### X-ray crystallography

Mutant GDAP1Δ303-358 crystals were obtained using sitting-drop vapour diffusion at +4 °C. Proteins were mixed with mother liquor on crystallisation plates using a Mosquito LCP nano-dispenser (TTP Labtech). The protein concentration was 10-30 mg/ml in 75 nl, and 150 nl of reservoir solution were added. R120Q crystals were obtained in 0.1 M succinic acid, 15% (w/v) PEG 3350. A247V crystals were obtained in 0.15 M *DL*-malic acid (pH 7.3), 20% (w/v) PEG3500. R282H crystals were obtained in 0.1 M succinic acid; 20% (w/v) PEG 3350. For cryoprotection, crystals were briefly soaked in a mixture containing 10% PEG200, 10% PEG400, and 30% glycerol, before flash cooling in liquid N2.

X-ray diffraction data were collected at the PETRA III synchrotron (DESY, Hamburg, Germany) on the P11 beamline [30, 31] and the EMBL/DESY P13 beamline [32] at 100 K and processed using XDS [33]. The structure of wild-type GDAP1Δ303-358, PDB entry 7ALM [11], was used as the search model for molecular replacement in Phaser [34]. The models were refined using Phenix.Refine [35] and rebuilt using COOT [36]. The structures were validated using MolProbity [37]. The data processing and structure refinement statistics are in **Table 1**, and the refined coordinates and structure factors were deposited at the Protein Data Bank with entry codes 7B2G (R120Q), 8A4J (A247V), and 8A4K (R282H).

**Table 1.** Data processing and refinement statistics. Data in parentheses refer to the highest-resolution shell.

| Variant | R120Q | A247V | R282H |
| --- | --- | --- | --- |
| Space group | P6<sub>3</sub>22 | P2<sub>1</sub>2<sub>1</sub>2<sub>1</sub> | P2<sub>1</sub>2<sub>1</sub>2<sub>1</sub> |
| Unit cell | 148.1, 148.1, 114.538 Å<br>90, 90, 120 ° | 73.3, 115.7, 115.9 Å<br>90, 90, 90 ° | 73.3, 113.4, 115.2 Å<br>90, 90, 90 ° |
| Resolution range (Å) | 100-3.0 (3.1-3.0) | 50-2.68 (2.84-2.68) | 50-1.95 (2.07-1.95) |
| Completeness (%) | 100 (100) | 99.7 (99.3) | 99.8 (99.9) |
| Redundancy | 38.7 (39.0) | 13.1 (13.5) | 6.7 (6.7) |
| $\langle I/\sigma \rangle$ | 17.0 (1.0) | 16.9 (1.9) | 15.4 (0.7) |
| R<sub>sym</sub> (%) | 27.3 (505.3) | 8.8 (162.7) | 4.9 (273.8) |
| R<sub>meas</sub> (%) | 27.7 (511.9) | 9.1 (169.0) | 5.3 (296.9) |
| CC<sub>1/2</sub> (%) | 99.9 (48.0) | 99.8 (81.4) | 99.9 (41.6) |
| R<sub>cryst</sub>/R<sub>free</sub> (%) | 23.0/26.5 | 25.2/28.3 | 22.5/25.1 |
| RMSD bond lengths (Å) | 0.002 | 0.015 | 0.014 |
| RMSD bond angles (°) | 0.5 | 1.5 | 1.4 |
| MolProbity score / percentile | 1.36 / 100<sup>th</sup> | 2.04 / 97<sup>th</sup> | 1.63 / 92<sup>nd</sup> |
| Ramachandran favoured/outliers (%) | 95.1/0.00 | 96.4 / 0.2 | 97.9 / 0.4 |
| PDB entry | 7B2G | 8A4J | 8A4K |

### Modelling

A model for full-length human GDAP1 was obtained from AlphaFold2 [38]. In addition, missing loops of the human wild-type GDAP1 crystal structure were built with CHARMM-GUI [39, 40]. The two models were used to shed light on conformational changes and sequence conservation in GDAP1. Earlier structure-based bioinformatics results [12] were analysed further with respect to the mutational spectrum of GDAP1.

### Synchrotron small-angle X-ray scattering

The structure and oligomeric state of the GDAP1 mutants were analysed with SEC-coupled small-angle X-ray scattering (SAXS). SEC-SAXS experiments were performed on the SWING beamline [41] (SOLEIL synchrotron, Saint Aubin, France). Samples were dialyzed against SEC buffer and centrifuged at >20000 g for 10 min at +4 °C to remove aggregates. 100 μl of each protein sample at 1.7-36 mg/ml were injected onto a BioSEC3-300 column (Agilent), run at a 0.3 ml/min flow rate. SAXS data were collected at +15 °C, over a q-range of 0.003–0.5 Å^-1^. SAXS data analysis, processing, and modelling were done in ATSAS 3.0 [42]. Scattering curves were analysed and particle dimensions determined using PRIMUS [43] and GNOM [44]. Chain-like *ab initio* models were generated using GASBOR [45], dummy atom models were built with DAMMIN [46], and model fitting to data was analysed with CRYSOL [47].

### Synchrotron radiation circular dichroism spectroscopy

Synchrotron radiation circular dichroism (SRCD) spectra were collected from 0.5 mg/ml samples on the AU-SRCD beamline at the ASTRID2 synchrotron (ISA, Aarhus, Denmark). The samples were prepared in a buffer containing 10 mM HEPES pH 7.5 and 100 mM NaF, equilibrated to room temperature, and applied into 0.1-mm closed circular quartz cuvettes (Suprasil, Hellma Analytics). SRCD spectra were recorded from 170 nm to 280 nm at +25 °C. Three scans per measurement were repeated and averaged. The spectra were processed using CDToolX [48].

### Thermal stability

Thermal stability of GDAP1 variants was studied by nanoDSF using a Prometheus NT.48 instrument (NanoTemper), in SEC buffer. Tryptophan fluorescence was excited at 280 nm, and emission was recorded at 330 and 350 nm, while the samples were heated from +20 to +90 °C at a rate of 1 °C/min. Changes in the fluorescence ratio (F350/F330) were used to determine apparent melting points. The data were analyzed using Origin (OriginLab Corporation, Northampton, MA, USA).

### Sequence entropy

Starting from the human GDAP1 reference sequence (NP_061845.2), iterative PSI-BLAST [49] searches were performed, initially accepting sequences with an e-value of 10^-99^ or smaller. From the first search, but the second iteration of PSI-BLAST, sequences were arbitrarily collected with e < 10^-07^. This resulted in 5986 sequences. This original data set was further subdivided into a first set that contains only sequences labeled GDAP1, a second set containing only sequences labeled GDAP1L1, and a third set containing only GST-labeled sequences. In this process, unnamed sequences, hypothetical sequences, and those from other proteins were discarded. Although GDAP1 and GDAP1L1 were initially separated, they will here be treated as one group (GDAP1/GDAP1L1). This is justifiable, since GDAP1L1 is by far the closest paralogue to GDAP1, and together they form an outlier group in the GST family. This was also a necessity, because the total number of sequences was too small compared to the GST data set. Even after merging the GDAP1/GDAP1L1 sets, there are still about twice as many sequences in the GST data set.

Lastly, all sets were combined to calculate the entropy across all data sets (GDAP1/GDAP1L1 + GST), which included 5065 sequences. For each set, a multiple sequence alignment was generated using MAFFT [50], and each sequence was mapped to a reference sequence. This was done by removing any columns, which correspond to a gap in the reference sequence. At this point, some information might get lost in the process. The first hit of human GDAP1L1 in the original data set was selected as reference. For the GST reference, a sequence was selected, for which there is a structure in the PDB database. This was not in the original data set and was added later (before alignment). **Supplementary Table 1** gives a detailed overview of the generated data sets.

For calculating the per-site entropy for a multiple sequence alignment, we used

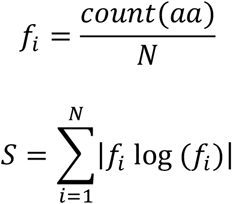

Where *f_i_* is the normalized frequency for each amino acid in each column of a MSA, and *N* is the number of sequences in the MSA, hence the number of rows. The basis of the logarithm depends on the alphabet (number of different symbols) and if gaps are treated as valid characters or should be ignored. For example, for protein sequences, the size of the alphabet is 20 without gaps and 21 with gaps. The entropy score S ranges from 0 (only one residue is present at that position) to 1 (all 20 residues are equally represented at the position). In this study, positions with S > 0.2 are considered variable, whereas those with S < 0.2 are considered conserved. Highly conserved positions are those with S < 0.1. Gaps are treated as non-valid characters.

### Kullback-Leibler divergence

The above chapter focused on the similarities between GDAP1/GDAP1L1 and GST. In addition to the similarities, differences are of particular importance. In order to capture these differences, the Kullback-Leibler (KL) divergence was used. The KL divergence is a statistical distance, a measure to quantify the differences between two probability distributions [51]. The KL divergence can be calculated as the sum of probability of each event in probability distribution P multiplied by the log of the probability of the event in probability distribution Q over the probability of the event in P [52]. Similarly to entropy calculation, the KL divergence score is derived from the normalized amino acid frequencies as probabilities:

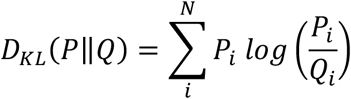

The intuition for the KL divergence score is that when the probability for an event from P is large, but the probability for the same event in Q is small, there is a large divergence. When the probability from P is small and the probability from Q is large, there is also a large divergence, but not as large as the first case. Importantly, the KL divergence score is asymmetrical:

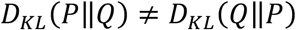

Hence, the KL divergence is calculated with one file as the P distribution and also as the Q distribution. In contrast to the above section, gaps are treated as valid characters. The goal is to find residues or regions that are highly conserved in both multiple sequence alignment files, but differ from each other. This should further shed light on the fundamental differences between GDAP1 and conventional GSTs.

### Phylogenetic analyses

To model the evolutionary history of GDAP1, the program MrBayes [53] was used. Visualizations were made with Interactive Tree of Life (iTOL) [54]. MrBayes is a Bayesian inference model that uses Markov chain Monte Carlo (MCMC) methods to estimate the posterior probability distribution of model parameters. Inference of phylogeny is based upon the posterior probability distribution of the constructed trees T_1_, T_2_,…, T_n_. The posterior probability of the i-th tree is calculated using Bayes’s theorem:

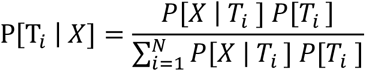

P[T_i_ | X] is the posterior probability of the i-th tree, P[X |T_i_] is the likelihood of the i-th tree, and P[T_i_] is the prior probability of the i-th tree. The denominator is a summation over all possible trees N. The likelihood P[X | T_i_] is calculated as a multidimensional integral over all possible combinations of branch lengths and substitution model.

The trees (with branch lengths) that were sampled by the MCMC procedure of each analysis were saved. The final tree is a consensus tree, an average over all calculated trees [55]. The settings used for the calculations are shown in **Table 2**. `Lset` is used to define the structure of the evolutionary model and `prset` is required to define the prior probabilities of the model. In this case (`prset aamodel=mixed`) the program integrates over a predetermined set of fixed rate matrices. `Ngen` sets the number of generations for which the analysis will be run. `Nruns` is the number of simultaneous, completely independent analyses starting from different random trees. `Nchains` is the number of chains used in each analysis. By setting `nchains=4`, MrBayes will use 3 heated chains and one “cold” chain. `Relburnin=yes` activates “burning”, which means that MrBayes will discard the first 25 % samples (`burninfrac=0.25`) from the cold chain. `Beagle` is a high-performance library for parallel computation.

**Table 2.** Parameter settings for MCMC analyses.

|  |
| --- |
| `prset aamodel= mixed` |
| `lset nucmodel=protein` |
| `Lset nst=1` |
| `Set autoclose=yes` |
| `Set usebeagle=yes` |
| `Set beagledevice=cpu` |
| `Mcmcp nruns=4 nchains =4` |
| `Mcmcp ngen= 2000000` |
| `Mcmp relburning= yes burninfrac=0.25` |
| `Mcmcp printfreq=1000000` |
| `Mcmcp savebrlens= yes` |

For phylogenetic analyses, PSI-BLAST was run as above, and from the 5986 sequences, GDAP1 sequences were extracted, resulting in 799 sequences, and the first hit of human GDAP1L1 (NP_001243666.1) was determined as the reference sequence for the GDAP1L1 set, and the first bacterial GST (TNE50161.1) was set as the bacterial GST reference sequence. Additionally, from the bacterial GST BLAST search, a sequence with a low e-value was set as the root sequence (WP_173192278.1). Since the BLAST search quickly arrives at bacterial GSTs, but it is also of interest where eukaryotic GSTs are found in the phylogenetic history, a human GST (PDB: 1PKZ), for which there is a crystal structure available, was set as a reference for eukaryotic GST. With each of these reference sequences (GDAP1L1, bacterial GST, eukaryotic GST), another BLAST search was performed, and the first 1000 hits of each BLAST search were extracted.

To study the relation of GDAP1 to archaea, a BLAST search with the GDAP1 reference was performed, but the output was restricted to archaeal sequences. With the first hit, an archaeal GST (MAE98075.1), another BLAST search was performed and the first 1000 hits were extracted, defining the last dataset (second_GST_set). Table 3 shows an overview over the separated datasets. Bacterial GST and eukaryotic GSTs were combined into the “first_GST_set”, to avoid oversampling. For each dataset, a multiple sequence alignment (MSA) was generated using MAFFT [Katoh et al., 2002]. Each set was then reduced to 100 sequences with the *reduce* program (*https://github.com/andrew-torda/sequtils*). *Reduce* generates an evenly distributed set of sequences by scanning the distance matrix and discarding very closely related sequences. The remaining 100 sequences of each dataset were combined, possible duplicated sequences were removed, and finally, together with the root sequence, re-aligned.

**Table 3.**
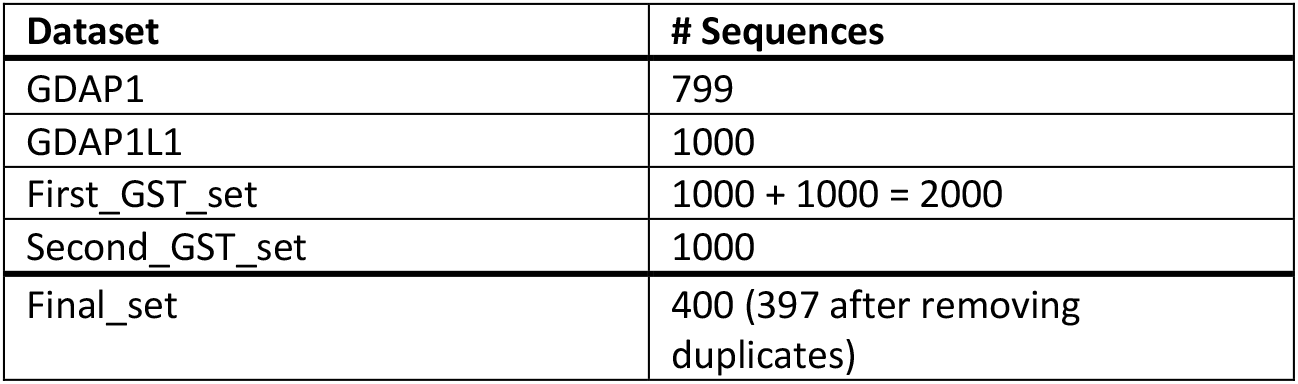
Dataset overview.

| Dataset | # Sequences |
| --- | --- |
| GDAP1 | 799 |
| GDAP1L1 | 1000 |
| First_GST_set | 1000 + 1000 = 2000 |
| Second_GST_set | 1000 |
| Final_set | 400 (397 after removing<br>duplicates) |

**Table 4.** SAXS parameters. Data for wild-type GDAP1 as well as the monomeric mutation Y29EC88E are taken from [11]. The MW estimate corresponds to the Bayesian estimate from PRIMUS.

| Variant | $\Delta 303$-358<br>dimer/<br>Y29EC88E<br>monomer | R120Q | R161H | A247V | H256R | R282H | R310Q<br>main peak | R310W<br>main peak | $\Delta 319$-358<br>dimer/mon<br>omer |
| --- | --- | --- | --- | --- | --- | --- | --- | --- | --- |
| $R_g$ (Guinier)<br>(Å) | 30.7/24.5 | 30.9 | 33.4 | 29.8 | 30.3 | 29.3 | 33.3 | 29.3 | 34.7/27.1 |
| $R_g$ (GNOM)<br>(Å) | 30.6/24.59 | 30.7 | 33.6 | 30.0 | 30.3 | 29.4 | 33.6 | 29.9 | 33.5/27.3 |
| $D_{max}$ (Å) | 99/86.7 | 99.7 | 110 | 100 | 99.2 | 91.0 | 120 | 110 | 107.9/89.6 |
| $V_p$ (nm<sup>3</sup>) | 105.8/58.7 | 102.0 | 127.4 | 93.1 | 95.8 | 89.4 | 101.2 | 58.2 | 129.5/69.4 |
| Estimated<br>MW (kDa) | 72.4/35.4 | 67.1 | 91.2 | 62.4 | 62.4 | 59.5 | 94.2 | 58.2 | 94.2/46.7 |

**Table 5.** nanoDSF apparent melting points. All values are average from 3 replicates.

| Sample | T<sub>m</sub> (°C) |
| --- | --- |
| GDAP1Δ303-358 | 62.6 |
| R120Q | 61.3 |
| R161H | 61.8 |
| A247V | 50.2 |
| H256R | 54.6 |
| R282H | 50.8 |
| GDAP1Δ319-358 | 56.9 |
| R310Q | 57.9 |
| R310W | 57.8 |

## RESULTS

Building upon earlier work on GDAP1 structure [10–12], we focused here on several CMT-linked variants that reside on different secondary structure elements on the GDAP1 crystal structure. While we earlier specifically looked at R120W and H123R on helix α3 [12], here we produced and characterised the variants R120Q, R161H, A247V, H256R, R282H, R310Q, and R310W. The stability and solution structure were studied for all variants, while crystal structures were determined for three of them: R120Q, A247V, and R282H. The location of the studied mutation sites in the wild-type GDAP1 structure is shown in **Fig. 1A-B**.

**Figure 1.**
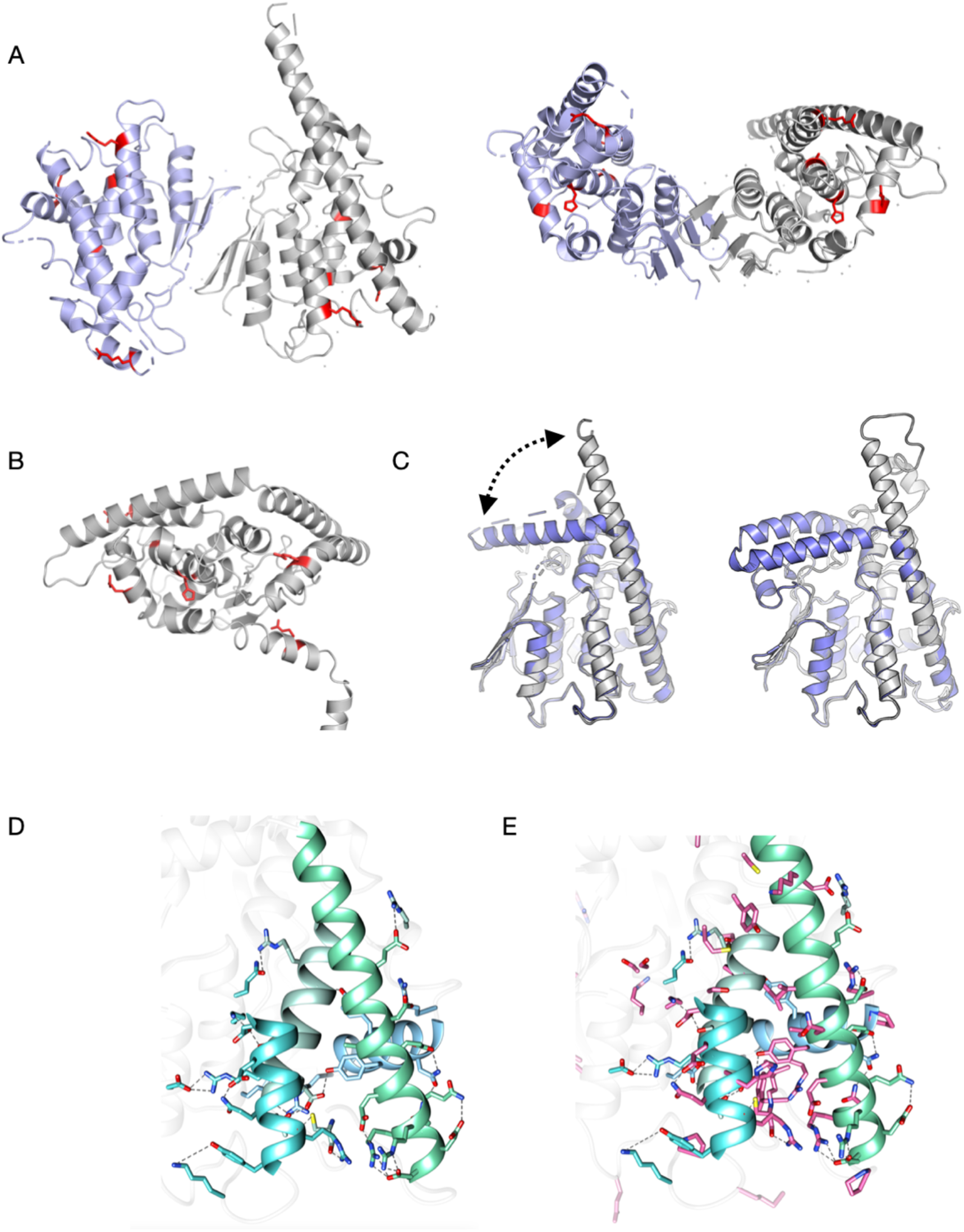
Overall structure of GDAP1. A. Location of the mutations studied here, mapped onto the crystal structure of wild-type GDAP1 [11] in two different orientations. B. The mapping of the mutations onto the AlphaFold2 model, to include those not visible in the crystallised construct (R161H, R310Q, R310W). C. Open/close conformations involving the long helix α6 have now been observed both experimentally (left) and using structure prediction (right). D. Hydrogen bonding network of residues on the core helices of GDAP1. E. Same view as E, but known sites of CMT mutations have been added in magenta. CMT mutations are clustered on the core helices.

### Helices α3, α6, α7, and α8 form the basis of GDAP1 intramolecular networks

The majority of CMT-linked missense mutations in GDAP1 are located within the vicinity of the hydrophobic clusters of the GST-like domains and the dimer interface [11], and the variants may induce changes in intramolecular hydrogen bonding networks [12]. In addition to the R120W and H123R studied earlier [12], we now determined three new mutant crystal structures: R120Q, A247V, and R282H. These mutations reside in helices α3 (R120Q), α7 (A247V), and α8 (R282H), which are core elements of the GDAP1 fold. We shall first look at the central helices regarding GDAP1 folding.

The GST-like core fold of GDAP1 is supported by the α7 helix, surrounded by helices α3, α6, and α8. The helix α3 is connected to α6 *via* the α6-α7 loop, and Cys240 in this loop - itself being a CMT mutation site - is central to many interactions. The α6 helix can either extend or turn back towards the dimer interface, as seen in earlier crystal structures and models [12]. A new model built here indicates that the extended conformation is predictable (**Fig. 1C**). Open/close movements of α6 can be functionally relevant for GDAP1 interactions with other proteins, such as cofilin and tubulin [9]. The α8 helix is positioned perpendicular to the others, and its orientation could be related to the transmembrane helix position, as it is expected to face the membrane surface. Together, these helices form an intramolecular network of polar (**Fig. 1D**) and non-polar contacts, and many CMT-linked missense mutations are found on these helices (**Fig. 1E).**

There are 15 designated hydrogen bond contacts or ionic interactions between helices α3, α6, α7, and α8. Helix α7, which is central to the GDAP1 fold, makes only a few hydrogen bonds to the surrounding helices. This indicates that hydrophobic residue clustering houses the α7 helix in the core of the fold; this is in line with centrality analyses of the GDAP1 fold, showing helix α7 to be the most central part of the 3D structure [12]. Some of the disease mutations on these helices correspond to solvent-exposed residues. Accordingly, helices α3, α6, and α8 show more polar interactions and higher flexibility compared to α7.

### Structural effects of the individual CMT mutations

Earlier, we analysed the effects of R120W and H123R, both on helix α3 [12]. Here, we extend the crystal structure analyses to three more CMT mutations: R120Q, A247V, and R282H (**Fig. 2, Table 1**).

**Figure 2.**
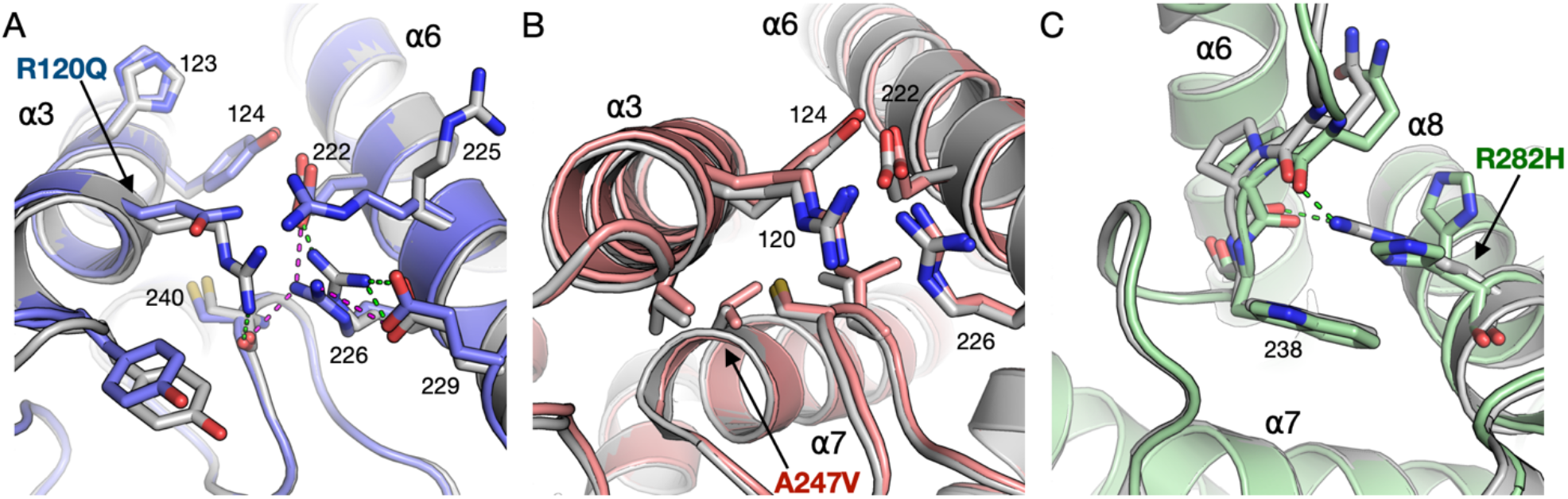
Structural details for each point mutation. A. Immediate environment of the R120Q mutation. B. Effects of the A247V mutation. C. The R282H mutation.

The R120Q mutation is located close to the N-terminal end of the α3 helix. Due to the mutation, residue 120 loses contact with the α6-α7 loop tip at Cys240, whereby the hydrogen bond between the Arg120 side chain and the backbone carbonyl of Cys240 is lost (**Fig. 2A**). This is further linked to alterations in surrounding side chain conformations.

The A247V mutation site resides at the N terminus of helix α7, tightly surrounded by helix α3 and the α6-α7 loop and having van der Waals contacts to Val121, Tyr124, Cys240, and Thr245. Comparing to the wild-type protein, changes in the crystal structure are minimal, but an overall movement of surrounding protein segments is caused by the presence of Val in this position. Both helices α3 and α6 move slightly away, without altering hydrogen-bonding patterns (**Fig. 2B**).

R282H is located on helix α8, and the side chain of Arg282 points inwards in the wild-type GDAP1 structure; it is stacked against Trp238 and makes hydrogen bonds to the backbone carbonyl groups of residues 236 and 237 in the α6-α7 loop (**Fig. 2C**). All these interactions are lost upon the mutation, and the His residue in the mutant is observed in a double conformation, making no side-chain hydrogen bond contacts.

### Conformation and stability in solution

Computational predictions suggested an overall destabilising effect of CMT-linked mutations in the GDAP1 protein [12]. Similarly, protein destabilisation was observed experimentally for the myelin protein P2 in the context of all identified CMT mutations [56, 57]. A comparative analysis of seven GDAP1 variants in solution was therefore carried out here, to support the crystal structures and other complementary data from the current and earlier studies, and to identify general trends linking CMT mutations and GDAP1 stability *in vitro.* While the 3D shape and dimensions were studied using SAXS, SRCD was employed to follow secondary structure content and nanoDSF to compare thermal stability.

The SAXS analysis of GDAP1Δ303-358 showed that the protein particle dimensions in solution correspond to a dimer, and the scattering profiles showed only minor shape differences, ruling out large-scale conformational differences or aggregation (**Fig. 3A**). The largest deviation was observed for R161H, which had a larger R_g_ than the other variants, suggesting a more open structure. Also R120Q had a slightly different conformation, most clearly seen in the distance distribution. The pair distance distribution function showed that the maximum particle dimension in all samples was ~90-110 Å, which corresponds to a dimer (**Fig. 3B**). *Ab initio* modelling, here carried out on the R282H mutant, which was measured at the highest concentration, agreed with the presence of a dimeric GDAP1, corresponding to the conformation of the dimer we previously showed to fit the solution SAXS data for wild-type GDAP1 [12] (**Fig. 3C**).

**Figure 3.**
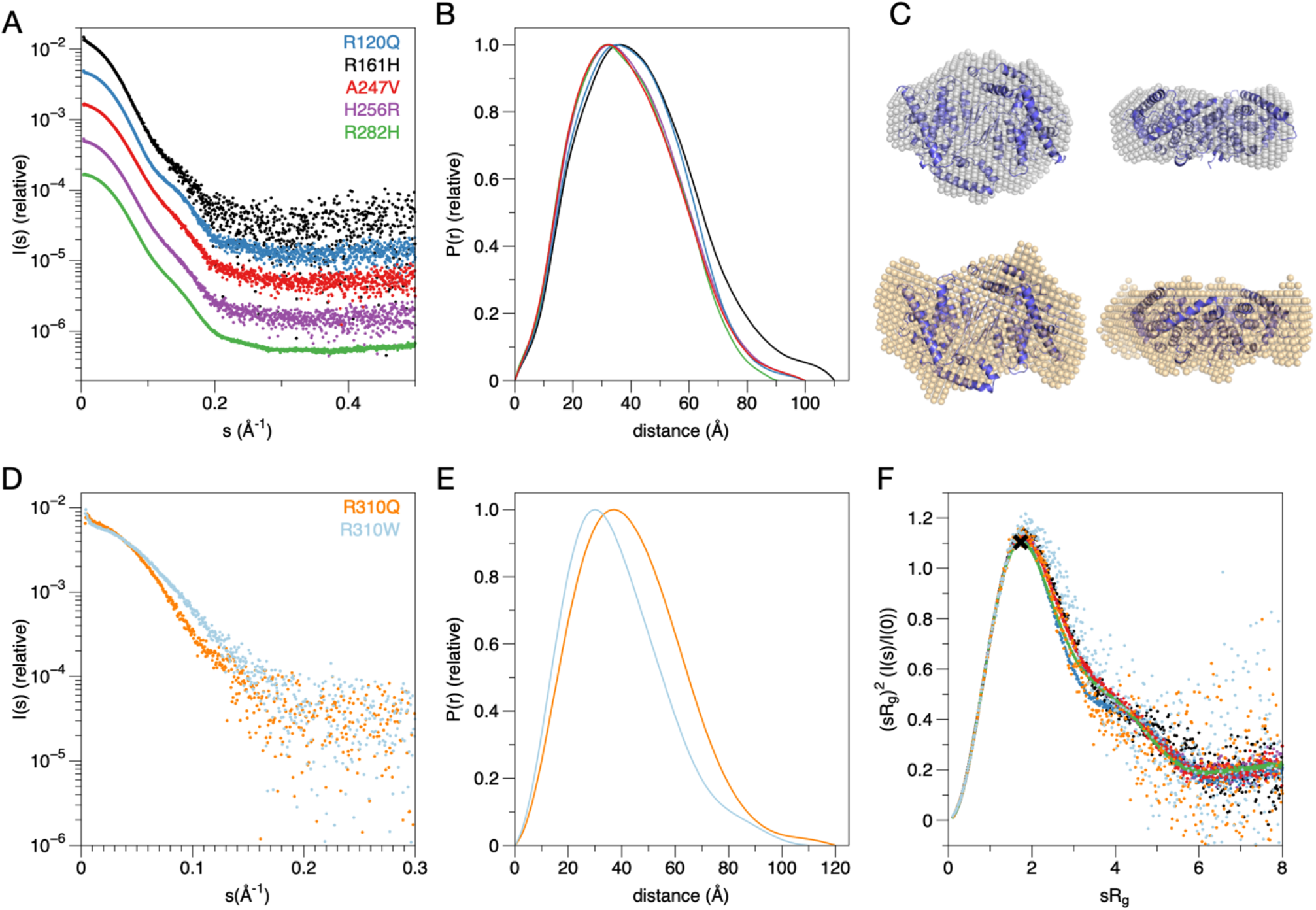
SAXS analysis. A. SAXS curves for the mutants in the GDAP1Δ303-358 construct. B. Distance distributions for the curves in A. C. Dummy atom model of R282H superimposed with a structure based on the collapsed conformations of wild-type GDAP1Δ295-358 [12] (top). Dummy atom model of the R310Q mutant in GDAP1Δ319-358, superimposed on the same structure. D. SAXS curves for the mutants in the GDAP1Δ319-358 construct. E. Distance distributions for the curves in D. F. Dimensionless Kratky plots for all constructs show similar levels of rigidity and globularity.

For the two mutations affecting Arg310, we used the longer construct GDAP1Δ319-358, and both R310Q and R310W eluted as two peaks in SEC-SAXS, corresponding to a dimer and monomer (**Fig. 3D-E**). This behaviour has previously been linked to protein concentration, in that monomers can be observed at low concentration [11], and the Arg310 mutants were here studied at lower concentration than all the other mutants. R310Q mainly eluted as a dimer, while the highest peak for R310W was monomer; since the protein concentrations were very similar, this result indicates that R310W may affect GDAP1 dimerisation. The R310Q dimer was more elongated than R282H of the shorter construct (**Fig. 3C**), reflecting the presence of the C-terminal amphipathic domain in the construct. Comparing the Kratky plot for all variants studied here, it is evident that they all are rigidly folded, with little flexibility (**Fig. 3F**).

For thermal stability and secondary structure analysis, nanoDSF and SRCD were performed. The SRCD spectra showed nearly identical secondary structure composition for all variants and wild-type GDAP1. Thus, none of the studied mutations affect the overall folding of GDAP1 (**Fig. 4A-B**). The DSF experiment revealed a ~1-12 °C decrease in apparent melting temperature for all core domain mutants, compared to the wild-type GDAP1Δ303-358 (**Fig. 4C**). This is similar to the previously studied R120W and H123R mutants [12]. All in all, the biophysical data indicate that the structural effects of all studied CMT variants in the core domain are local, and the mutations do not large-scale conformational changes at a level detectable with SRCD or SAXS. However, all studied mutations in the core domain caused a decrease in thermal stability, suggesting a breakdown of stabilising intramolecular interactions within the GDAP1 molecule. As the most dramatic example, the apparently harmless mutation A247V decreased the stability by 12.4 °C.

**Figure 4.**
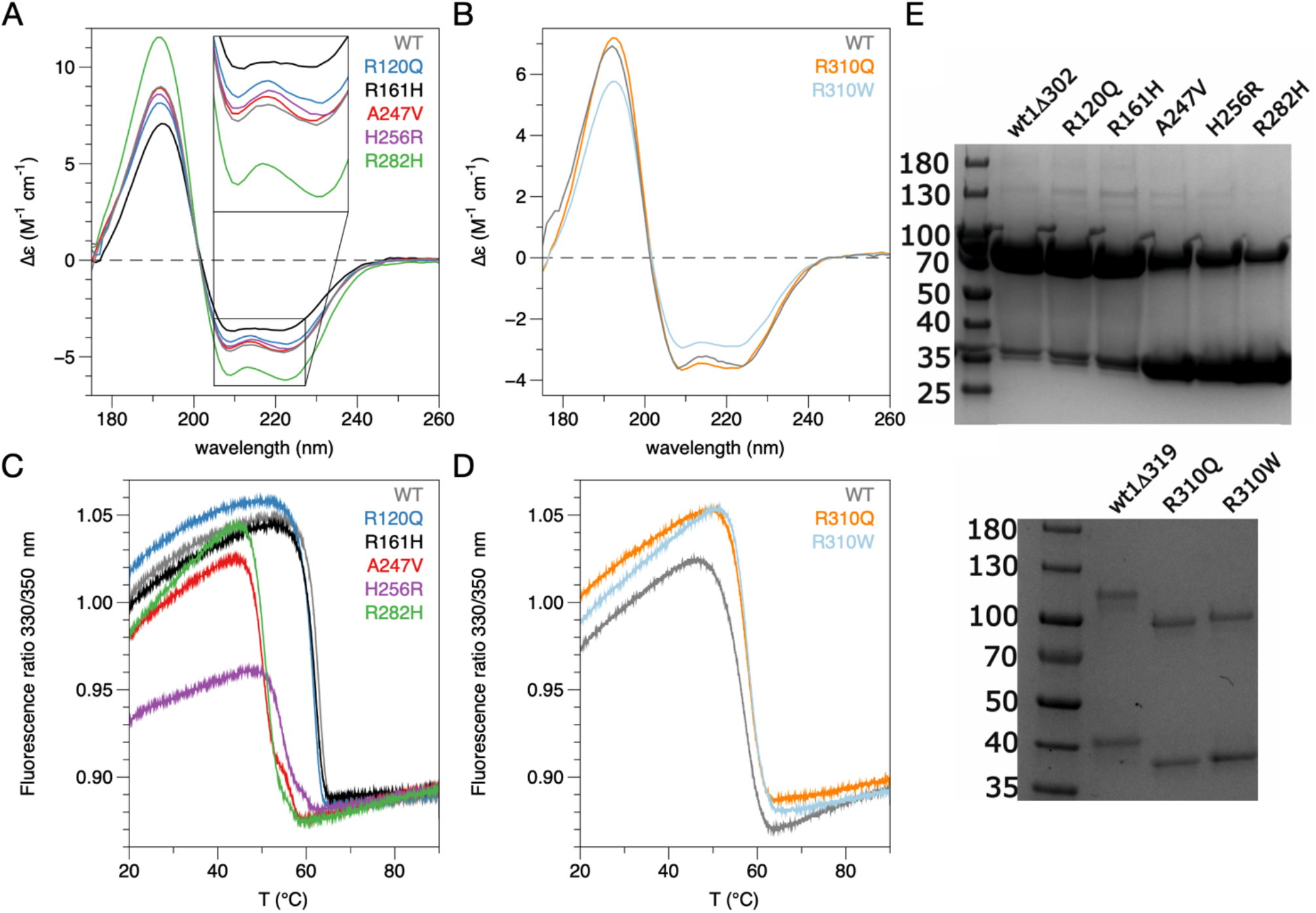
Folding and stability. A,B. nanoDSF analysis for the mutants in GDAP1Δ303-358 (A) and GDAP1Δ319-358 (B). C. SRCD spectra for the mutants in the GDAP1Δ303-358 construct. D. SRCD spectra for the mutants in GDAP1Δ319-358. E. SDS-PAGE analysis of all studied variants.

For the mutations at position 310, it was observed that both R310Q and R310W in fact increased the stability of GDAP1Δ319-358 by ~1 °C (**Fig. 4D**). This can be explained by the fact that the segment carrying these mutations is no longer part of the core domain, but rather likely to represent an *a* helix attached to the membrane surface, possibly linking the membrane thus to the core domain. Overall, one can say GDAP1 stability correlates with the dimer-monomer ratio observed in solution; the most destabilised mutants also show a higher fraction of monomeric protein on non-reducing SDS-PAGE (**Fig. 4E**), even though in the 3D structure, they are far from the dimer interface.

### Implications of the point mutations for the whole protein

The studied mutations cause subtle variations in hydrogen bond and van der Waals distances in nearby residues in the crystal structures, while there are no drastic structural differences, when the mutant structures are superposed to the wild-type crystal structure. However, when comparing the variation of the residues in helices α3, α6, α7, and α8, in comparison to the wild-type Cα-atoms, a specific pattern arises. We analysed the residues participating a hydrogen-binding network in the mutant structures (**Fig. 5A**), and the mapping shows that the residues pointing towards the dimer interface have only minor variations compared to the wild-type protein. In contrast, at the C-terminal end of helix α6, in all mutant crystal structures, the variation of the Cα atoms is high (**Fig. 5B**). This suggests that flexibility of helix α6 could arise from altered intramolecular contacts in the vicinity. The C-terminal end of the α6 helix, the conformation of which apparently is affected by the mutations studied here, is itself a target for multiple CMT mutations affecting Gln218, Va219, Glu222, Arg226, and Glu227 [19, 58–61].

**Figure 5.**
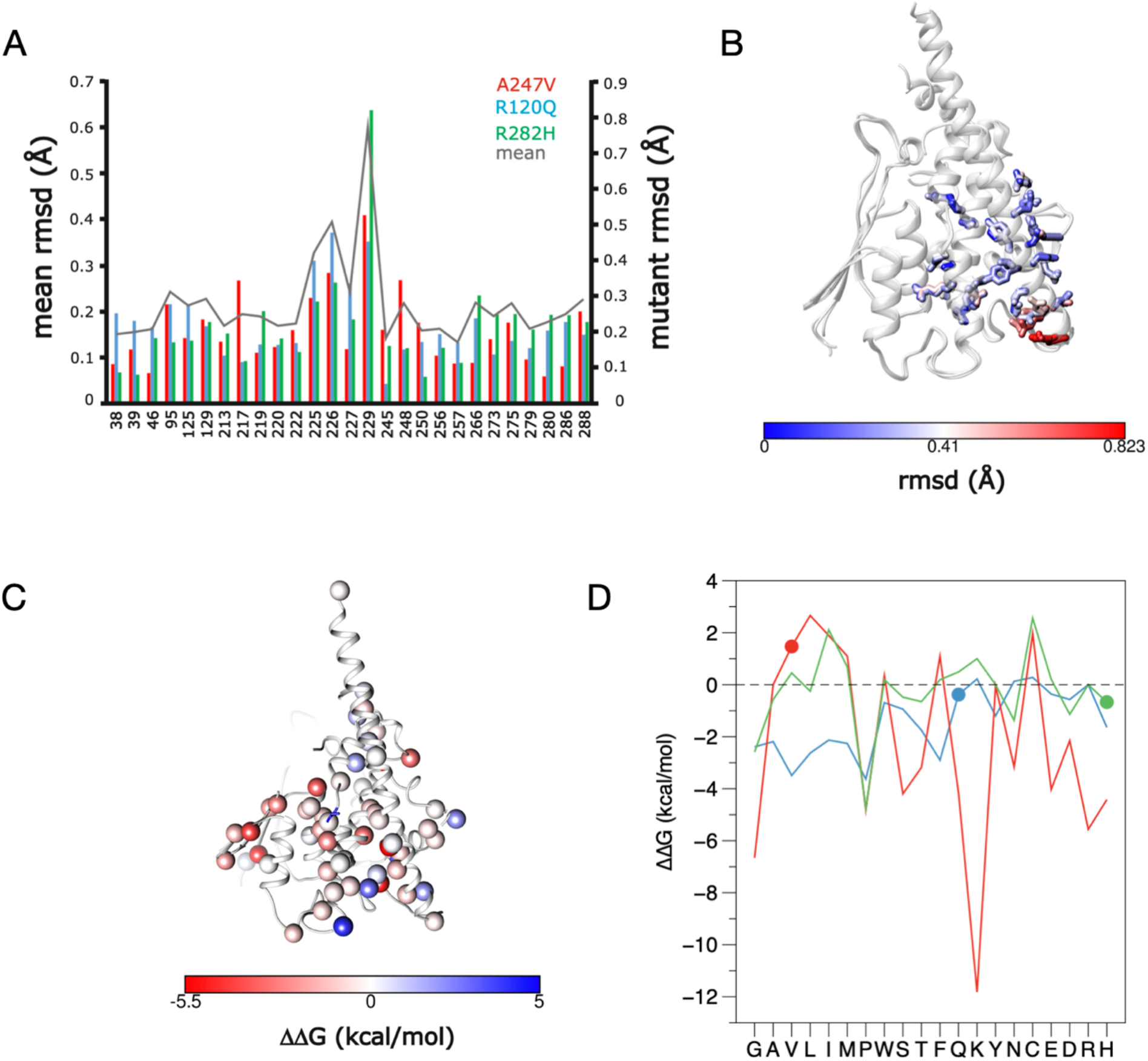
Structural bioinformatics. A. The Ca deviation of each mutant vs. wild-type GDAP1 structure. Shown are only the residues participating in the hydrogen bonding network. B. Mapping the results onto the structure, it becomes evident that the C-terminal end of helix α6 deviates the most from wild-type GDAP1 on average. C. Average ΔΔG effect of mutations at CMT sites, as defined by CUPSAT. D. CUPSAT predictions for R120Q, A247V, and R282H being mutated into all possible amino acids. Note that a negative ΔΔG in CUPSAT means destabilisation. The mutations studied here are marked with spheres.

We additionally investigated the more global mutational effects using bioinformatics tools. In our previous study [12], we analysed the wild-type GDAP1 core domain structure and the full-length AlphaFold2 coordinates with CUPSAT and MAESTRO [62, 63]. This analysis provided predicted ΔΔG values and geometry properties; the results from CUPSAT analyses are further depicted in **Fig. 5C**. Clearly, on the average most CMT mutation sites are predicted to cause destabilisation and the few residues predicted to be stabilised by mutations lie on the protein surface.

While the above analyses indicate an average effect that may destabilise GDAP1 structure, we looked at predictions for each mutation studied here in more detail (**Fig. 5D**). In essence, all substitutions to Arg120 are unfavourable; this shows that the interactions made by Arg120 are important for folding and stability. This is in line with our experimental data for both R120Q and R120W, which indicate minor changes in structure, but destabilisation of the fold. For Ala247, a mutation into Val is actually predicted to be slightly stabilising, indicating that such a replacement, in a tightly confined pocket and with small but long-range effects, is difficult for the prediction algorithm. In the case of Arg282, mutation to His is indeed one of the most destabilising variants in the prediction.

### Phylogenetic analysis of GDAP1 and the GST superfamily

Since GSTs are universally widespread across almost all organisms, the GST superfamily is inherently very large and diverse. There are massive differences between families and even within the subfamilies. Starting a BLAST search with the reference sequence will quickly end up in GSTs. This leads to the fact that GSTs are oversampled compared to GDAP1 and GDAP1L1; respectively, GDAP1/GDAP1L1 are undersampled. Therefore, the phylogenetic tree calculates the evolutionary history of GSTs, rather than the evolutionary history of GDAP1. To avoid this, GDAP1, GDAP1L1 and GSTs were considered separately.

As a first step in a sequence-based bioinformatics approach, we carried out a phylogenetic analysis of the GST family with a specific focus on GDAP1 (**Fig. 6**). Intriguingly, based on large-scale sequence alignments, GDAP1 turns out to be more closely related to prokaryotic GSTs than eukaryotic proteins. In this situation, we must remember that sequence conservation levels in the GST superfamily are overall low. Based on the phylogenetic analysis, it is still not possible to depict the exact pathways of GDAP1 evolution. In general terms, one can say GDAP1 is at least as far from generic eukaryotic GSTs as it is from prokaryotic proteins. The analysis provides little clues as to any enzymatic function of GDAP1, but it does give more support to the findings that it has no conventional GST activity.

**Figure 6.**
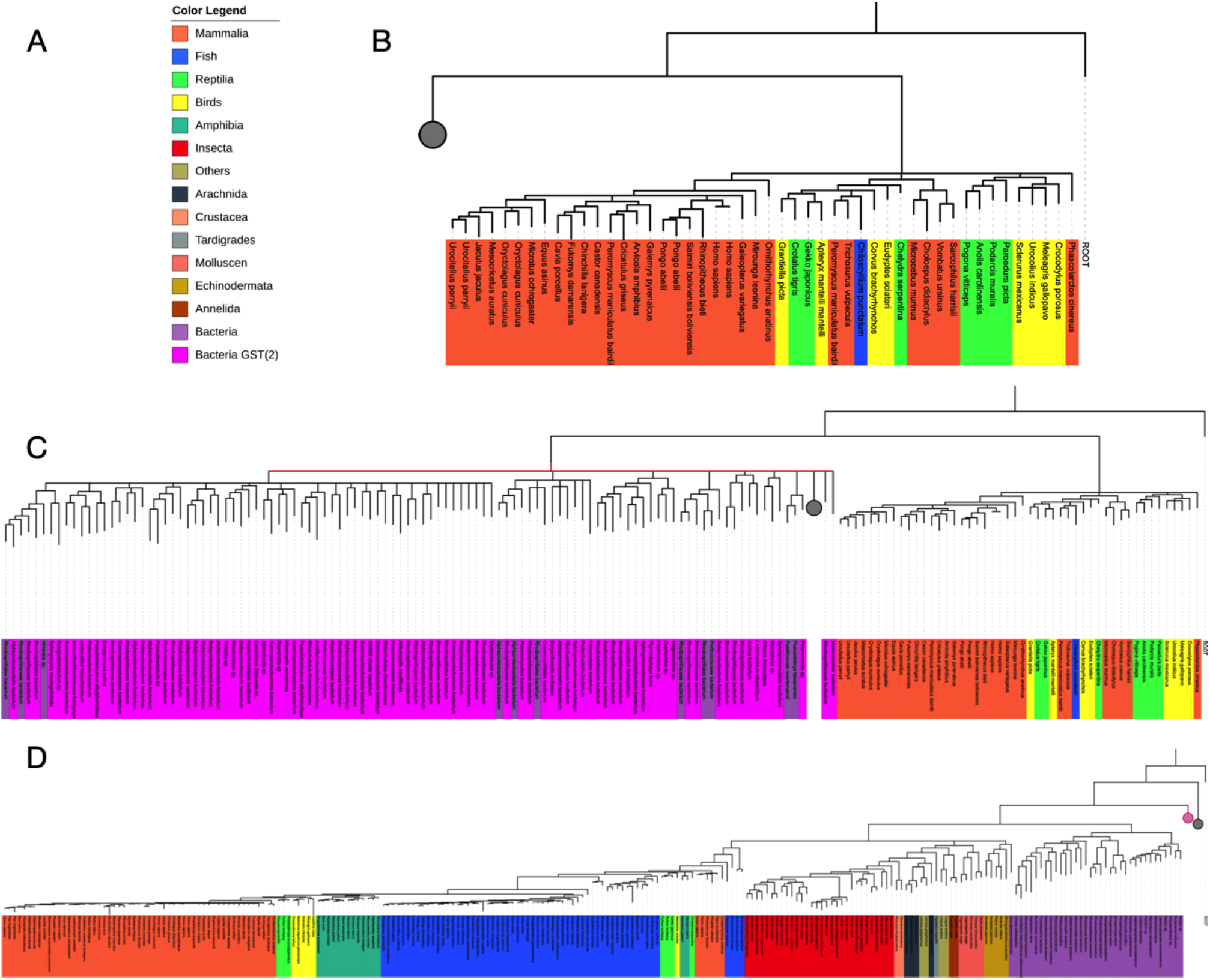
Phylogenetic analyses. A. Colour scheme for phylogenetic trees. Note that not all organisms seen in the colour legend are present in every picture. This is because the trees are very large and to increase visibility parts of the tree will be collapsed (denoted by a grey circle), to hide underlying branches. B. The first branch of the phylogenetic tree. The tree was rooted at the root sequence. On the right side are eukaryotic GST sequences (mammals, fish, birds, reptiles). The branch includes 48 sequences. The rest of the 398 sequences that are represented are at hierarchically lower levels of the phylogenetic tree. The corresponding branches were collapsed (grey circle). C. In contrast to panel B, the left branch is partially extended. The right-sided branch shows eukaryotic GST sequences, while the left side shows a total of 110 bacterial GST sequences. The remaining sequences are at hierarchically lower levels of the phylogenetic tree. The corresponding branches were collapsed (grey circle). The red line indicates that the time could not be resolved at this level. D. The last hierarchical level of the phylogenetic tree. The tree was rooted at the root sequence. Purple sequences on the right side correspond to bacterial GST sequences. Sequence for eukaryotic GDAP1 and GDAP1L1 are in the left side branches. The grey circle denotes the collapsed branch for the eukaryotic GSTs (see panel B) and the magenta circle denotes the second group of bacterial GST sequences (see panel C).

**Fig. 6B and 6C** show that eukaryotic GSTs split off from other GSTs very early in evolutionary history. This was not to be expected. In addition, **Fig. 6B** shows that the evolutionary history at this level could not be resolved (represented by a red line) despite exhausting CPU time (**Table 2**). One can see that it is a bifurcating tree. Bacterial GST sequences are shown in the right branch and eukaryotic GST sequences in the left branch. The remaining sequences in the tree are represented by the grey circle.

**Fig. 6C** shows the last level of the phylogenetic tree. Considering only this part, this tree is a bifurcating tree. Towards the right side there is a branching to bacterial GST sequences (purple) and in the left branching the GDAP1/GDAP1L1 sequences are found again. The figure also shows that orthologs of GDAP1 and GDAP1L1 are found in mammals, birds, amphibians, and fishes and likely orthologs of those genes in invertebrates as well.

A first assumption of the evolutionary history is based on the endosymbiotic theory and the current perception of the phylogenetic tree of life that archaea are evolutionarily closer to eukaryotes than bacteria [64]. Since GDAP1 is a protein of the outer mitochondrial membrane, it is possible that GDAP1 is closely related to archaeal proteins. Since the BLAST search with an archaeal sequence almost exclusively finds bacterial GST sequences, and the fact that the times could not be fully resolved, this tree does not make any assumptions whether GDAP1 are derived from archaea or not. However, it does suggest early multiple gene duplication events that split up eukaryotic GST sequences.

### Sequence entropy in GDAP1

The entropy analysis was carried out on both the whole GST family, only the GDAP1/GDAP1L1 sequences, or only on non-GDAP1 sequences. We will first look at the results from the GDAP1 subset of sequences (**Fig. 7A**). A total of 44 residues were found with an entropy score ≤ 0.10 and a percentage of non-gap characters present in the alignment > 70 (**Table 6**); the entropy is mapped onto the GDAP1 structure in **Fig. 7B**. The data highlight several interesting residues, which are known to be sites of CMT mutations, relevant for folding and structure, or with possible functions in ligand binding. These aspects are discussed more below.

**Figure 7.**
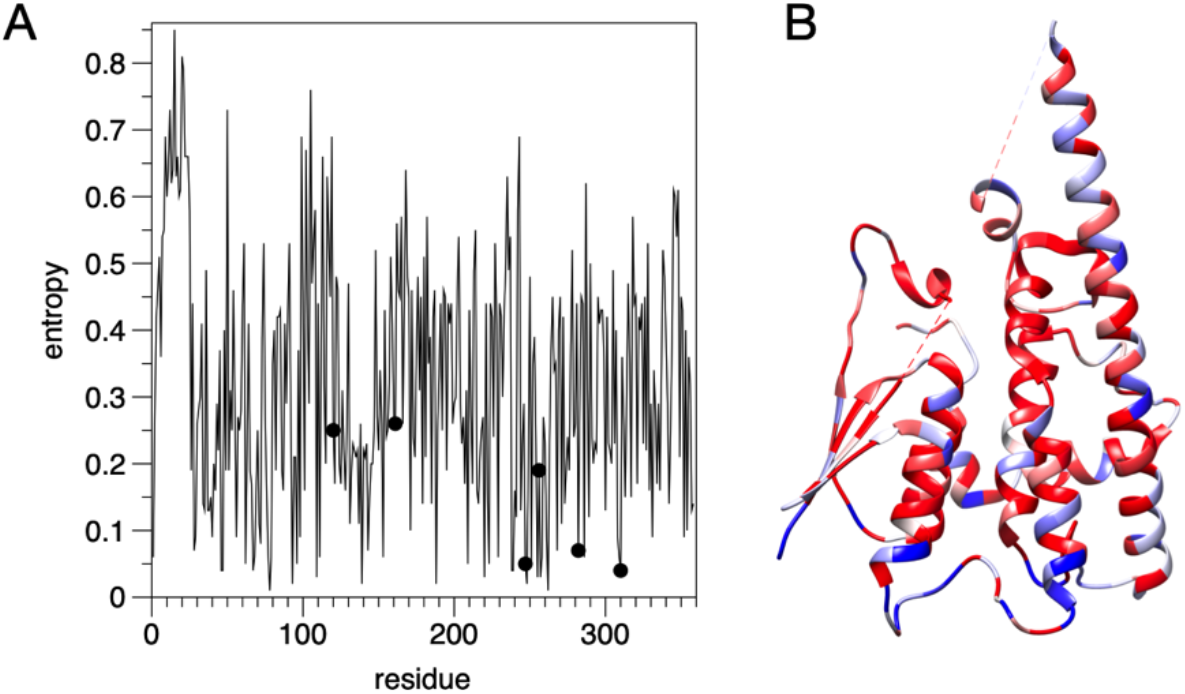
Sequence entropy analysis. A. Entropy plot for GDAP1. The positions of the mutations studied here are shown with dots. B. Mapping of entropy onto the GDAP1 crystal structure monomer. Blue indicates high entropy and red low.

**Table 6.** Entropy analysis of the GDAP1 subfamily. Shown are the positions (human GDAP1 reference sequence numbering) with the lowest entropy (S < 0.1).

| residu<br>e | entrop<br>y | frac | Amino<br>acid | residu<br>e | entrop<br>y | frac | Amino<br>acid | residu<br>e | entrop<br>y | frac | Amino<br>acid |
| --- | --- | --- | --- | --- | --- | --- | --- | --- | --- | --- | --- |
| 28 | 0.07 | 0.78 | L | 98 | 0.09 | 0.94 | E | 248 | 0.02 | 0.97 | D |
| 29 | 0.09 | 0.78 | Y | 109 | 0.03 | 0.87 | L | 251 | 0.05 | 0.97 | L |
| 40 | 0.09 | 0.83 | V | 111 | 0.06 | 0.91 | P | 255 | 0.03 | 0.97 | L |
| 46 | 0.04 | 0.83 | E | 139 | 0.02 | 0.91 | G | 257 | 0.03 | 0.97 | R |
| 47 | 0.04 | 0.83 | K | 143 | 0.07 | 0.91 | H | 258 | 0.06 | 0.97 | L |
| 56 | 0.09 | 0.83 | V | 153 | 0.06 | 0.91 | P | 261 | 0.09 | 0.97 | L |
| 62 | 0.05 | 0.84 | E | 171 | 0.1 | 0.94 | L | 262 | 0.01 | 0.96 | G |
| 67 | 0.04 | 0.84 | W | 188 | 0.02 | 0.94 | K | 268 | 0.07 | 0.96 | W |
| 68 | 0.06 | 0.84 | F | 209 | 0.05 | 0.94 | L | 282 | 0.07 | 0.96 | R |
| 72 | 0.08 | 0.93 | N | 220 | 0.03 | 0.94 | E | 286 | 0.06 | 0.96 | R |
| 77 | 0.06 | 0.93 | V | 223 | 0.05 | 0.94 | L | 308 | 0.09 | 0.84 | A |
| 78 | 0.01 | 0.93 | P | 229 | 0.06 | 0.82 | E | 309 | 0.06 | 0.82 | F |
| 79 | 0.06 | 0.93 | V | 238 | 0.04 | 0.97 | W | 310 | 0.04 | 0.82 | R |
| 93 | 0.02 | 0.93 | I | 239 | 0.04 | 0.97 | L | 331 | 0.09 | 0.89 | G |
| 96 | 0.05 | 0.94 | Y | 247 | 0.05 | 0.97 | A |  |  |  |  |

One intuitive approach to search for functionally important sites is to focus on residues involved in ligand binding, *i.e.* residues that are located in the active centre. However, the question of whether GDAP1 is an active GST, or even lacks glutathione binding completely, has not been clearly resolved. Hexadecanedioic acid was identified as possible ligand and subsequently crystallized in complex with GDAP1 [11]. Gln235, Trp238, Arg282, Arg286, and Lys287 contact the ligand; Trp238, Arg282 and Arg286 are highly conserved (**Table 6**). This suggests that they could be important for ligand binding and that mutations in these positions could affect GDAP1 function; indeed, Trp238 and Arg282 are CMT mutation sites. These 3 residues cluster on the side of GDAP1, which must face the membrane surface. One option is that the bound fatty acid, in fact, mimics the membrane surface, and that these residues are directly involved in membrane surface binding at the outer surface of the MOM.

**Table 7** shows the results of conservation entropy based on the combined data set (GDAP1, GDAP1L1, and GST). Since the GST superfamily is diverse by nature, with a conserved fold but low sequence conservation, the limits were relaxed. All residues with an entropy ≤ 0.20 and a percentage of non-gap character > 70 are listed in the table. Overall, 14 residues were derived by entropy calculations. Note that these include Leu239 and Gly241 in the α6-α7 loop and Ala247 and Asp 248 on helix α7. The effect of the A247V mutation on protein stability may therefore be linked to its high conservation in the GST family.

**Table 7.** Entropy analysis for the large data set including the GST dataset. Shown are residue positions with the lowest entropy (S < 0.2).

| residue | entropy | frac | Amino acid |
| --- | --- | --- | --- |
| 28 | 0.12 | 0.92 | L |
| 72 | 0.15 | 0.98 | N |
| 77 | 0.17 | 0.98 | V |
| 78 | 0.01 | 0.98 | P |
| 93 | 0.05 | 0.98 | I |
| 96 | 0.1 | 0.98 | Y |
| 109 | 0.17 | 0.9 | L |
| 111 | 0.14 | 0.81 | P |
| 223 | 0.1 | 0.89 | L |
| 239 | 0.2 | 0.9 | L |
| 241 | 0.14 | 0.9 | G |
| 247 | 0.12 | 0.9 | A |
| 248 | 0.02 | 0.9 | D |
| 286 | 0.17 | 0.86 | R |

### Kullback-Leibler divergence

The results of the KL divergence are shown in **Table 8**. Many listed residues are insertions because they correspond to a gap symbol in the alignment. Of the GDAP1 residues Cys51, Gly83, Tyr124, and Glu228 that are not assigned to a gap symbol, Cys51 and Gly83 are located close to the dimerization site of GDAP1. Gly83 is a target of a non-pathological polymorphism, G83A [65]. The β sheet is the dimerization site of GDAP1, and the larger amino acids Phe and Asp may lead to steric hindrance during dimer formation. On the other hand, Phe and Asp are predestined for stacking and polar interactions. Since the binding site for glutathione in GST takes place near this segment, these residues could be important for ligand binding in the GSTs, which may not to be relevant for GDAP1 due to the apparent lack of glutathione binding. The observations support the assumption that the binding site for a small-molecule ligand, if any, in GDAP1 may be located elsewhere.

**Table 8.** KL divergence.

| res num | kIP | kIQ | Sp | Sq | GDAP1 | GST |
| --- | --- | --- | --- | --- | --- | --- |
| 51 | 1.0292121 | 0.776726 | 0.1934639 | 0.46464223 | C | F |
| 83 | 0.711117 | 0.8625605 | 0.4245107 | 0.37385303 | G | D |
| 106 | 1.608235 | 1.8079219 | 0.47457752 | 0.26693898 | T | - |
| 124 | 1.1740065 | 1.2500156 | 0.19621734 | 0.38821948 | Y | W |
| 131 | 0.9714733 | 2.912868 | 0.11059844 | 0.16480081 | L | - |
| 132 | 0.94019866 | 1.8085836 | 0.19164778 | 0.39762115 | P | - |
| 149 | 0.7277371 | 2.1168306 | 0.27086747 | 0.26017815 | D | - |
| 150 | 0.703265 | 1.8003652 | 0.21712388 | 0.2821153 | S | - |
| 151 | 1.8435297 | 2.2269633 | 0.3448432 | 0.3719185 | M | - |
| 166 | 1.2843894 | 2.3159494 | 0.2552226 | 0.25735173 | N | - |
| 186 | 1.1256604 | 2.4295797 | 0.41708586 | 0.2931384 | I | - |
| 192 | 1.0507396 | 1.881601 | 0.24847703 | 0.4722129 | L | - |
| 193 | 0.86935514 | 2.3180199 | 0.4641341 | 0.4705446 | K | - |
| 228 | 1.1552744 | 1.9867854 | 0.40772098 | 0.3858474 | E | Q |
| 229 | 1.6037626 | 1.9517808 | 0.06304588 | 0.48370922 | E | - |
| 233 | 2.1693683 | 2.274921 | 0.42683157 | 0.312741 | E | - |
| 234 | 2.1514575 | 2.6264071 | 0.48738602 | -0.0 | G | - |
| 236 | 2.176767 | 2.9161973 | 0.48846936 | -0.0 | Q | - |
| 295 | 1.5544128 | 2.371196 | 0.45351568 | 0.49149024 | H | - |
| 300 | 2.4423957 | 2.5055954 | 0.12804487 | 0.4074487 | L | - |
| 319 | 1.0532062 | 2.999142 | 0.4438014 | 0.21979499 | V | - |
| 339 | 1.537214 | 1.3012601 | 0.48322365 | 0.352141 | L | - |
| 352 | 2.5170794 | 1.6161157 | 0.094565034 | 0.42042798 | R | - |
| 353 | 1.3202384 | 1.7585322 | 0.39914683 | 0.4678713 | P | - |
| 355 | 2.271911 | 1.898613 | 0.3553678 | -0.0 | P | - |
| 356 | 2.2988195 | 1.9953156 | 0.32429653 | 0.41148844 | N | - |
| 358 | 2.5016084 | 2.5750391 | 0.1393856 | -0.0 | F | - |

On the other hand, Tyr124 and Glu228 are close to each other in 3D space, being located on the core helices α3 and α6, respectively. The conversion of tyrosine to tryptophan (Tyr124 → Trp) and glutamic acid to glutamine (Glu228 → Gln) between GDAP1 and canonical GSTs causes only small changes in physicochemical properties. However, the central location of these residues in the GDAP1 fold suggests this finding may reflect a structural or functional aspect of GDAP1.

Of additional interest are Lys193 and Asn166, which are both close to the GDAP1-specific insertion. In contrast to the crystal structure, where Asn166 is not visible, the AlphaFold2 model predicts that the helix α6 is bent, which brings Lys193 and Asn166 into close proximity. This could enable intramolecular interactions or provide a specific site for protein-protein interactions. Overall, the results indicate that certain subdomains in GDAP1 may have evolved to fulfill a different function than found in canonical GSTs.

## DISCUSSION

Mutations in dozens of genes are causative for various subtypes of CMT disease; for most of them, effects of missense mutations at the protein level remain poorly characterised. This is partly because the biological functions of the corresponding proteins are poorly understood at the molecular level. One of the characterised examples is myelin protein P2, which loses thermal stability upon all the known 6 CMT mutations in the protein, while the crystal structure remains nearly unaltered [56, 57]. Furthermore, the disordered tail of myelin protein P0 is a target for P0 mutations, and its membrane interactions may be compromised upon CMT [66]. Periaxin carries several CMT-linked truncation mutations that abolish protein-protein interactions [67, 68], most notably with β4 integrin [69]. While these proteins, highly enriched in myelinating Schwann cells, are involved in the classical Schwann cell phenotypes related to myelination, it is evident that compromised mitochondrial function is one underlying cause of especially axonal CMT subtypes, and mutations in GDAP1 are linked to mitochondrial dysfunction. Linked to this mechanism, recent data show that GDAP1 may be involved in interactions with the actin and tubulin cytoskeletons [9].

Mutations in the GST-like domain of GDAP1 have a broad pathological spectrum. The molecular basis is still unknown despite the cellular observations confirming the causality of impaired mitochondrial dynamics, and accurate structural information is required to support these findings. Below, we shall discuss some implications of our findings to understand GDAP1 function at the molecular level and the effects of missense mutations therein.

### CMT mutations at the GDAP1 protein level

CMT-linked missense mutations are relatively common in the *GDAP1* gene compared to other CMT target genes, especially when the size of the protein is taken into consideration [23, 70]. With careful examination of structural models and using them as inputs for further bioinformatics analyses, the mutations are observed to cause subtle changes in residue interaction networks, in line with experimental data. A comprehensive understanding of the effects of single mutations requires local observation of the structure coupled to experimental data at the protein and cellular levels.

Of the residues highlighted by the entropy analysis either within the GDAP1 sequence set or the entire GST set, several are targets for missense mutations. Here, we shall briefly compare selected CMT mutations in GDAP1. The immediate environment of each residue is considered to shed light on local effects of each mutation.

Tyr29 is conserved at the symmetry axis of the GDAP1 dimer, forming a H bond between OH groups of Tyr29 from the two protomers. Y29S [71] would both remove this polar interaction and make the dimer interface much less hydrophobic.

Leu239 is at the tip of the α6-α7 loop, inserted into the structural core, and its mutation to Phe has been reported in CMT [72]. The side chain is close to those of Cys240 and Ala247, and hence, a larger residue at this position could similarly affect protein stability as A247V. Similar effects could be foreseen for the C240Y mutation [73]. Pro111 is in the a2-a3 loop, close in 3D space to the N terminus of helix α7 and the α6-α7 loop. Its mutation to His [74] would cause steric hindrance and alter the conformation of the a2-a3 loop, possibly destabilising the fold.

Ala247, which was studied in the current work, lies centrally in the GDAP1 fold on helix α7. A247V is linked to CMT, showing that no larger residue fits into this tight space. A247V caused destabilisation of the GDAP1 fold. Ala247 is one of the most conserved residues in the GST superfamily, indicating a role in the GST fold. Other residues conserved in the GDAP1 set include Leu255 and Gly262, which lie in the middle and at the end of helix α7, respectively. L255F [75] would be expected to cause similar steric hindrance as A247V, and G262E [71] will likely disturb the tight turn right after helix α7 and cause steric clashes. Both Leu255 and Gly262 are highly conserved in GDAP1 sequences.

Arg282, studied here in the form of the CMT mutation R282H, is one of the most conserved residues in the GDAP1/GDAP1L1 subfamily. Its strong hydrogen bonding to the carbonyl groups in the α6-α7 loop indicates a central role in GDAP1 structure. In addition to R282H, the mutation R282C has been reported [76], and we can expect it to have the same kind of effect as R282H, losing the side-chain interactions of Arg282 to the backbone of the α6-α7 loop. Arg282 is stacked against the aromatic side chain of Trp238, which is both as a conserved residue and a CMT mutation target [77]. This Arg-Trp-(α6-α7) unit is likely important for stable folding of GDAP1.

Arg310 is not present in the constructs we used for crystallisation, and hence, we have no high-resolution data on its conformation. However, the AF2 model of GDAP1 extends our crystal structures and predicts that Arg310 points away from the membrane and forms two salt bridges to acidic residues in the GDAP1 fold (**Fig. 8**). Interestingly, Arg310 resides in the segment originally coined the hydrophobic domain; in our view, this fragment more likely corresponds to an amphipathic helix, which could bind onto a phospholipid membrane surface. The AlphaFold2 prediction supports this view. Arg310 is likely to link this membrane-bound helix to the folded core of GDAP1 and therefore directly affect the conformation of full-length, membrane-bound GDAP1. This is supported by the lack of a destabilising effect on recombinant GDAP1 structure in our experiments.

**Figure 8.**
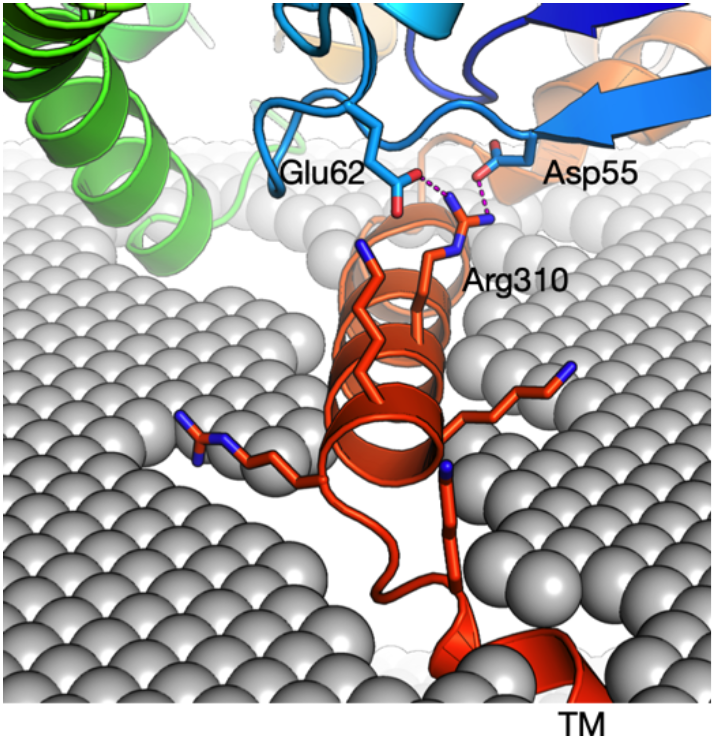
Location of Arg310 outside the GST-like core. Arg310 within the amphipathic helix preceding the transmembrane domain is predicted to make salt bridges back towards the core domain.

The clinical profiling of CMT patients and experiments in cell-based models have shown that residues located within or near the vicinity of the transmembrane helix are severe [78, 79]. These mutations likely affect the proper localization into the MOM, leading to impaired GDAP1 folding and function. So far, studies *in vitro* on the GDAP1 protein have been done only on soluble constructs, lacking the transmembrane domain. The stability analysis of the mutant form of the full-length GDAP1 could have a different outcome due to the lipid bilayer component, and this is a future research direction, given that full-length recombinant GDAP1 can be expressed and purified.

### The central role of the α6-α7 loop

Despite the broad spectrum of disease mutations, certain general conclusions hold in terms of the mutations. The interaction network described here shows that the critical helices in the C-terminal GST-like core are linked by residues that correspond to CMT target locations and/or are conserved. Therefore, based on evolutionary reasoning, these residues are likely essential for the integrity of GDAP1 and its function.

The α6-α7 loop is a central feature in the structure of GDAP1, but also in canonical GSTs. This loop inserts itself back into the protein, being a central interaction hub between helices α3, α6, α7, and α8 (**Fig. 9**). In GDAP1, the 3 residues at the tip of the loop (238-240) are targets for CMT disease mutations, showing the crucial importance of this segment. Our entropy analysis confirms the strong conservation of this segment. We can then take a broader scope and further look at residues interacting with the α6-α7 loop in GDAP1. Of note, several of the residues in direct contact with the α6-α7 loop are known CMT mutation target sites and/or highly conserved (**Table 6**). Hence, the inward-bending α6-α7 loop appears to be a central structural feature in GDAP1, and its alteration either directly or *via* intramolecular interactions may be a common mechanism for CMT.

**Figure 9.**
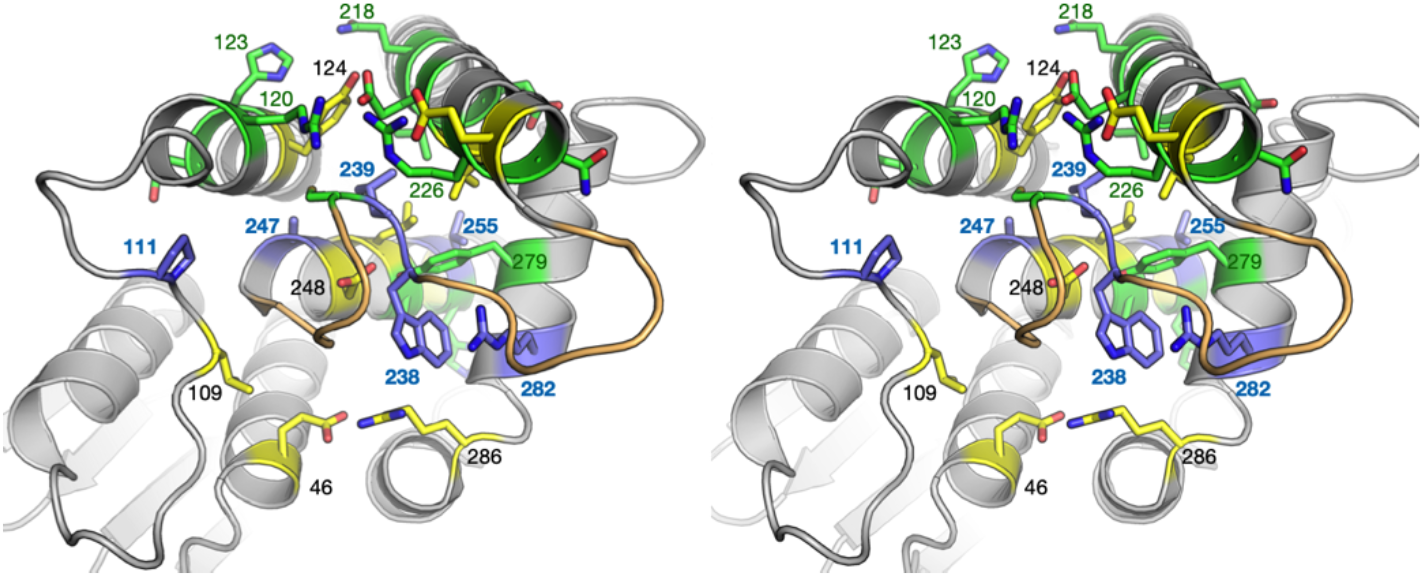
An overview of the α6-α7 loop and its surroundings in light of CMT mutations and conserved positions. In this stereo view, the α6-α7 loop is coloured orange, green shows positions for CMT mutations, yellow positions that are highly conserved, and blue the positions that are both targeted by CMT mutations and highly conserved. Selected residues are labelled for clarity.

### GDAP1 as a member of the GST superfamily

The predictions that GDAP1 would be structurally related to GSTs have been confirmed by recent structural studies by us and others [10–12]. Important differences to canonical GSTs exist, however. The sequence identity to any enzyme with known GST activity is very low, and contradictory results have been obtained as to the GST activity of GDAP1. In our hands, GDAP1 does not bind glutathione or act as a GST [11]. The latter is logical, since the GDAP1 dimers are formed completely differently from canonical GST, in which the active site in fact lies at the dimer interface [80]. We hypothesise that the residues with low entropy and high conservation in the whole GST set are crucial for correct folding. On the other hand, residues additionally highlighted in the GDAP1/GDAP1L1 dataset may relate to GDAP1-specific functions and/or unique structural aspects of this subfamily.

Our conservation analyses highlight the low conservation of GDAP1 (and GDAP1L1) in the GST superfamily. While the entire evolutionary pathway cannot be tracked based on the analyses, it is intriguing that GDAP1 is closer to bacterial than eukaryotic GSTs. It is possible that during evolution, GDAP1 has lost the characteristic GST activity, while becoming an integral membrane protein of the MOM. Its functions could, therefore, be mediated through protein-protein interactions instead of enzymatic activity. The functions can be redox-regulated, which could explain the observation that the human GDAP1 dimer is mediated by a disulphide bridge *via* Cys88 [11].

To complement the above analyses, we superposed crystal structures of human GDAP1 and a canonical GST, that from *S. japonicum* [80], and analysed the current crystal structures with respect to GST. This is the GST widely used in molecular biology applications as a fusion tag for affinity purification. We were interested in the residues affected by CMT mutations, especially those crystallised here. Hence, of specific interest were the apparent non-conservation of Arg120 and the conservation of Ala247 and Arg282 (**Fig. 10A**).

**Figure 10.**
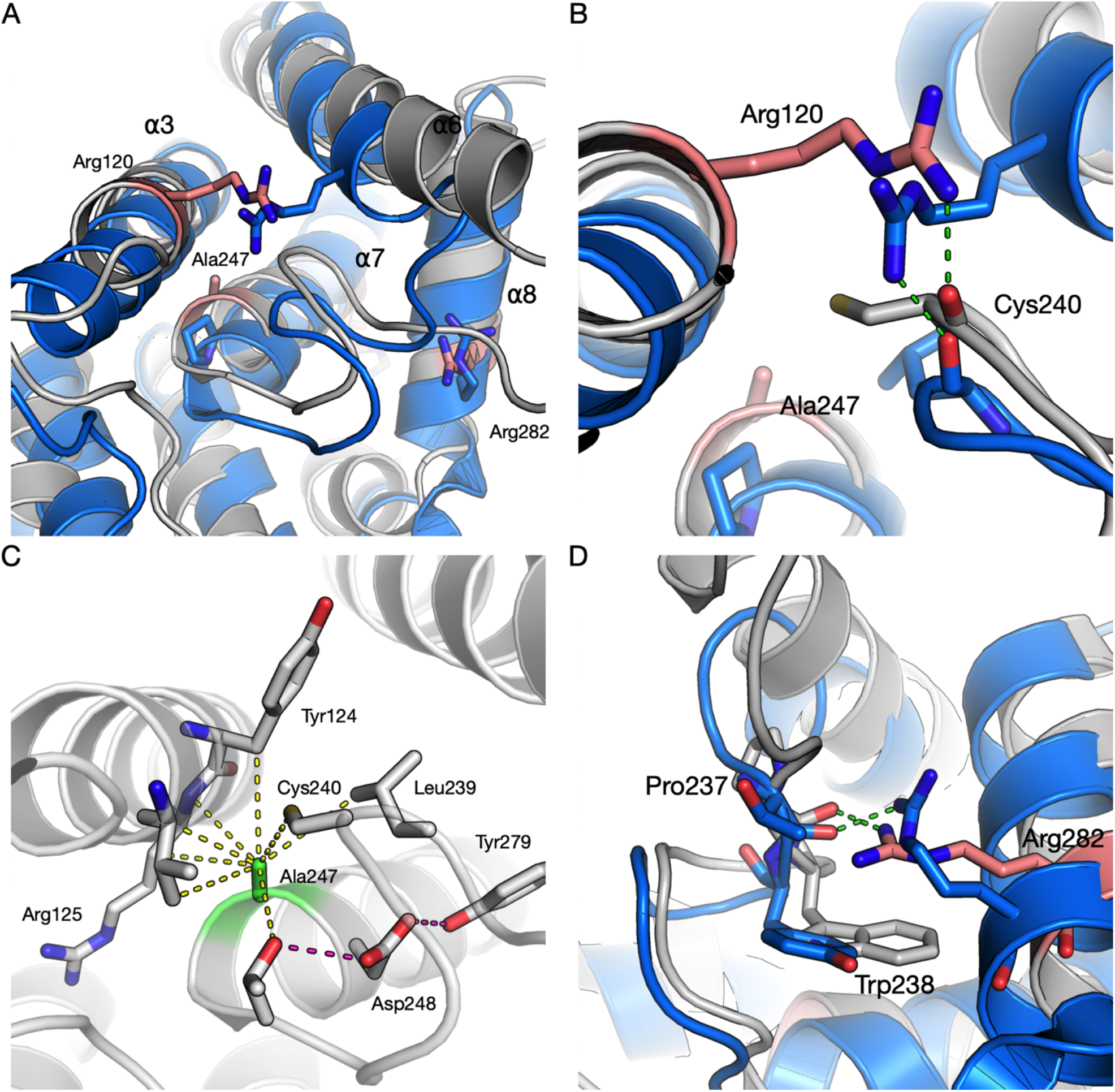
Comparisons of the studied mutations to a canonical GST from S. japonicum. GDAP1 is in gray and GST in blue. A. Overall view of the GDAP1-GST superposition, with the CMT mutation sites crystallised here highlighted. B. Arg120 (pink) in GDAP1 and the corresponding interaction in GST. C. Tight packing of Ala247 (green) in the GDAP1 structure; Ala at this position is highly conserved across the whole GST family. Note Tyr124, which was highlighted in the KL divergence analysis, making direct contact with Ala247. D. The interactions of Arg282 (pink) towards the α6-α7 loop are conserved in GST.

Arg120 in GDAP1 is a target of several CMT mutations [81–83], and its interactions with the backbone carbonyl of Cys240 in the α6-α7 loop appear central. Structures of the mutants R120Q and R120W indicate loosening of the structure locally, as well as loss of the hydrogen bond, which is accompanied by a decrease in observed heat stability in R120Q and R120W [12]. Arg120 is not conserved in GSTs at the sequence level; however, an Arg residue from the neighbouring helix in GST reaches to the same position and makes similar interactions to the same carbonyl group in the α6-α7 loop (**Fig. 10B**). These observations highlight the importance of the central α6-α7 loop in intramolecular interaction networks and fold stability of GDAP1.

Ala247 of GDAP1, although perhaps a mundane residue *per se*, appears surprisingly conserved in the GST superfamily in our dataset (**Table 6**). In the superposed individual GST structure (**Fig. 10A**), this residue is Pro, which fits well into the structure due to slightly different conformations of the main helices in GST. In GDAP1, Ala247 is so snugly packed (**Fig. 10C**) that even the addition of two methyl groups in the A247V variant causes protein instability and disease, despite minor effects in the crystal structure. A247V is an example of a mutation introduced into the hydrophobic core that may seem relatively harmless, but in fact has long-range effects on the entire protein structure stability. One of the residues in GDAP1 highlighted by the KL divergence analysis, Tyr124, closely interacts with Ala247, indicating a GDAP1-specific arrangement at this site. Additionally, one of the most conserved GDAP1 residues, Asp248, is central in a hydrogen bonding network linking the conserved CMT target Tyr279 into the picture (**Fig. 10C**).

GST has an Arg corresponding to Arg282, and this residue is highly conserved in the GST family, as shown by our bioinformatics analyses (**Table 7**). Arg282 in GDAP1 interacts directly with the backbone of the α6-α7 loop, and a similar interaction is observed in GST (**Fig. 10D**). This is another example of an Arg - loop backbone interaction conserved and relevant for human disease mutations.

### Biological implications

The function of GDAP1 at the molecular level remains enigmatic to date. While indications exist from functional studies [2, 3, 79], evidence is still incomplete for any enzymatic activity as well as direct protein-protein interactions. Redox regulation seems to play a role [9, 73, 84], but is this related to an enzymatic GST-like activity, or regulation of oligomeric state and/or protein-protein interactions?

At the molecular level, we believe to have identified important residue interaction networks between the core helices in the GST-like domain of GDAP1, strongly interacting with the inward α6-α7 loop. These networks could be important for both GDAP1 stability and its interactions with other molecules, such as the membrane or the cytoskeleton. The correct conformation of GDAP1 on the MOM, as well as its interactions with other proteins, will then directly or indirectly affect mitochondrial dynamics to promote correct development and function of the nervous system. The disease mutations may – due to their involvement in the same intramolecular networks – cause similar overall effects on GDAP1 stability and properties, which then leads to the CMT disease phenotype in patients. In line with this, all the missense mutations we have studied at the protein level allow for GDAP1 folding, while at the same time decreasing the stability of the fold.

It should be noted that due to the large pool of GDAP1 mutants causing CMT, our experimental sample size is still relatively small, and the hypothesis may not be correct in all cases. However, we have picked mutations from different core secondary structure elements for experimental analyses to account for an incomplete dataset, and predictions have been done for all mutations [12], indicating a general trend of structural destabilisation upon CMT missense mutations in GDAP1.

### Concluding remarks

Considering mitochondrial dynamics and interactions with other organelles, the homology to GSTs brings attractive prospects for GDAP1 function. GDAP1 takes part not only in mitochondrial fission and fusion, but in interactions with the endoplasmic reticulum, peroxisomes, Golgi, and the cytoskeleton. These findings coupled with structural and biophysical data shall aid the understanding of the pathophysiological mechanism of GDAP1-linked CMT and may affect future treatment approaches. Future studies are needed to identify proteins and small molecules directly interacting with full-length GDAP1 in the physiological setting, allowing further structural investigations on the related molecular processes in nervous system function and disease. In a wider setting, we hypothesise that decrease in overall protein stability upon missense mutations is one common mechanism for CMT at the molecular level, and many of the related interaction networks are linked to the core helices and the α6-α7 loop of GDAP1.

## Acknowledgements

This work was funded by the Academy of Finland, project number 24302881. We wish to acknowledge the availability of synchrotron beamtime and excellent beamline support at both SOLEIL, ISA, and DESY. The SAXS and SRCD experiments were supported by CalipsoPlus, which is funding from the European Union Horizon 2020 research and innovation programme under grant agreement No 730872. SRCD experiments further received funding from the European Union Horizon 2020 research and innovation programme under grant agreement No 101004806 (MOSBRI-2021-24).

